# Oxidized alkyl phospholipids stimulate sodium transport in proximal tubules via PPARγ-dependent pathway

**DOI:** 10.1101/2021.05.02.442360

**Authors:** Tomohito Mizuno, Nobuhiko Satoh, Shoko Horita, Hiroyuki Tsukada, Yusuke Sato, Haruki Kume, Masaomi Nangaku, Motonobu Nakamura

## Abstract

The pleiotropic effects of oxidized phospholipids (oxPLs) have been identified. 1-O-hexadecyl-2-azelaoyl-*sn*-glycero-3-phosphocholine (azPC), an oxPL formed from alkyl phosphatidylcholines, is a potent peroxisome proliferator-activated receptor γ (PPARγ) agonist. Although it has been reported that thiazolidinediones can induce volume expansion by enhancing renal sodium and water retention, the role of azPC, an endogenous PPARγ agonist, in renal transport functions is unknown. In the present study, we investigated the effect of azPC on renal proximal tubule (PT) transport using isolated PTs and kidney cortex tissues. We showed that azPC rapidly stimulated Na^+^/HCO_3_^-^ cotransporter 1 activity and luminal Na^+^/H^+^ exchanger (NHE) activities in a dose-dependent manner, at submicromolar concentrations, in isolated PTs from rats and humans. Additionally, the stimulatory effects were completely blocked by a specific PPARγ antagonist, 2-chloro-5-nitro-N-phenylbenzamide (GW9662), and a mitogen-activated protein/extracellular signal-regulated kinase (MEK) inhibitor, PD98059. Treatment with an siRNA against PPARγ significantly suppressed the expression of PPARγ mRNA, and it completely blocked the stimulation of both Na^+^/HCO_3_^-^ cotransporter 1 and NHE activities by azPC. Moreover, azPC induced extracellular signal-regulated kinase (ERK) phosphorylation in rat and human kidney cortex tissues, and the induced ERK phosphorylation by azPC was completely suppressed by GW9662 and PD98059. These results suggest that azPC stimulates renal PT sodium-coupled bicarbonate transport via the PPARγ/MEK/ERK pathway. The stimulatory effects of azPC on PT transport may be partially involved in the development of volume expansion.

## Introduction

Hypertension can lead to the development and progression of atherosclerosis, and it also contributes to the development of cardiovascular diseases (1). In contrast, atherosclerosis is known to be a risk factor for hypertension (2). However, the underlying mechanism for the development of hypertension due to atherosclerosis has not been elucidated yet. A large number of studies have demonstrated the role of oxidation products in the progression of atherosclerosis (3, 4). Due to oxidative stress, oxidized phospholipids (oxPLs) are generated from a variety of phospholipids containing polyunsaturated fatty acids (3). OxPLs are mainly accumulated in atherosclerotic lesions (5), and they are associated with endothelial dysfunction (3, 6), adhesion, transmigration, cytokine production by macrophages (5, 7), proliferation, migration, and phenotypic switching of vascular smooth muscle cells (VSMCs) (8–10), and apoptosis (7). Although oxPLs exert both proatherogenic and protective effects by affecting diverse gene expression and signaling pathways, their proatherogenic action is predominant at the sites of tissue deposition of oxPLs, leading to the progression of atherosclerosis (5,11,12).

An increase in renal proximal tubule (PT) sodium reabsorption can lead to hypertension (13). Approximately 50%-60% filtered Na^+^ and 80% filtered HCO_3_^-^ are reabsorbed from PT by the cooperative action of Na^+^/H^+^ exchanger 3 (NHE3) and vacuolar-type H^+^-ATPase (V-ATPase) expressed on the luminal membranes and Na^+^/HCO_3_ cotransporter 1 (NBCe1) expressed on the basolateral membranes (14–16). The functions of NHE3 and NBCe1 are regulated by humoral factors and various signaling mechanisms (14, 17). Indeed, we have previously reported some NHE3 and NBCe1 stimulators, such as angiotensin II (Ang II), insulin, and thiazolidinediones (TZDs) (18–22).

Peroxisome proliferator-activated receptor γ (PPARγ), a ligand-activated transcription factor belonging to the nuclear receptor superfamily, is expressed in various tissues and cell types, such as white and brown adipose tissues, VSMCs, macrophages, and vascular endothelial cells (23). PPARγ is also widely present in the kidney, including PTs and collecting ducts (24). TZDs are well-known exogenous PPARγ agonists that exert pleiotropic effects, including an improvement of insulin sensitivity and anti-inflammatory effects (25). The use of TZDs has been limited due to important side effects such as edema and congestive heart failure (26, 27). TZD-induced volume expansion is largely due to an enhancement of renal sodium and water retention (28). Additionally, TZDs stimulate both NBCe1 and NHE3 activities through the PPARγ/proto-oncogene tyrosine-protein kinase Src/epidermal growth factor receptor (EGFR)/extracellular signal-regulated kinase (ERK)-dependent nongenomic signaling pathway in isolated rat, rabbit, and human PTs (22). On the other hand, a variety of endogenous PPARγ ligands such as oxidized low-density lipoproteins (oxLDLs), oxPLs, eicosanoids, and linoleate derivatives have also been identified (29, 30). 1-O-hexadecyl-2-azelaoyl-*sn*-glycero-3-phosphocholine (azPC), an oxidation product of LDL alkyl phosphatidylcholines (PCs) present in atherosclerotic lesions, is a potent PPARγ agonist (30). The binding affinity of azPC is almost equivalent to that of rosiglitazone (30). However, the influence of azPC on renal sodium and fluid transport remains unclear.

Therefore, in the present study, we used isolated PTs from rats, mice, and humans to investigate whether azPC affects renal PT sodium transport and TZDs *in vitro* and *ex vivo*.

## Results

### Effects of azPC on NBCe1 activity in isolated rat PTs

To investigate the effects of azPC on PT transport, we first examined NBCe1 activity using freshly isolated and luminally collapsed PTs from rat kidneys. As shown in Figures 1A and S1, azPC rapidly stimulated NBCe1 activity in isolated rat PTs. The stimulatory effects of azPC on NBCe1 activity were dose-dependent in the concentration range 0.04 to 0.3 μM while no difference was observed between 0.3 μM azPC and 1.0 μM azPC. Therefore, we conducted further experiments using 0.3 μM azPC.

**Figure 1.**
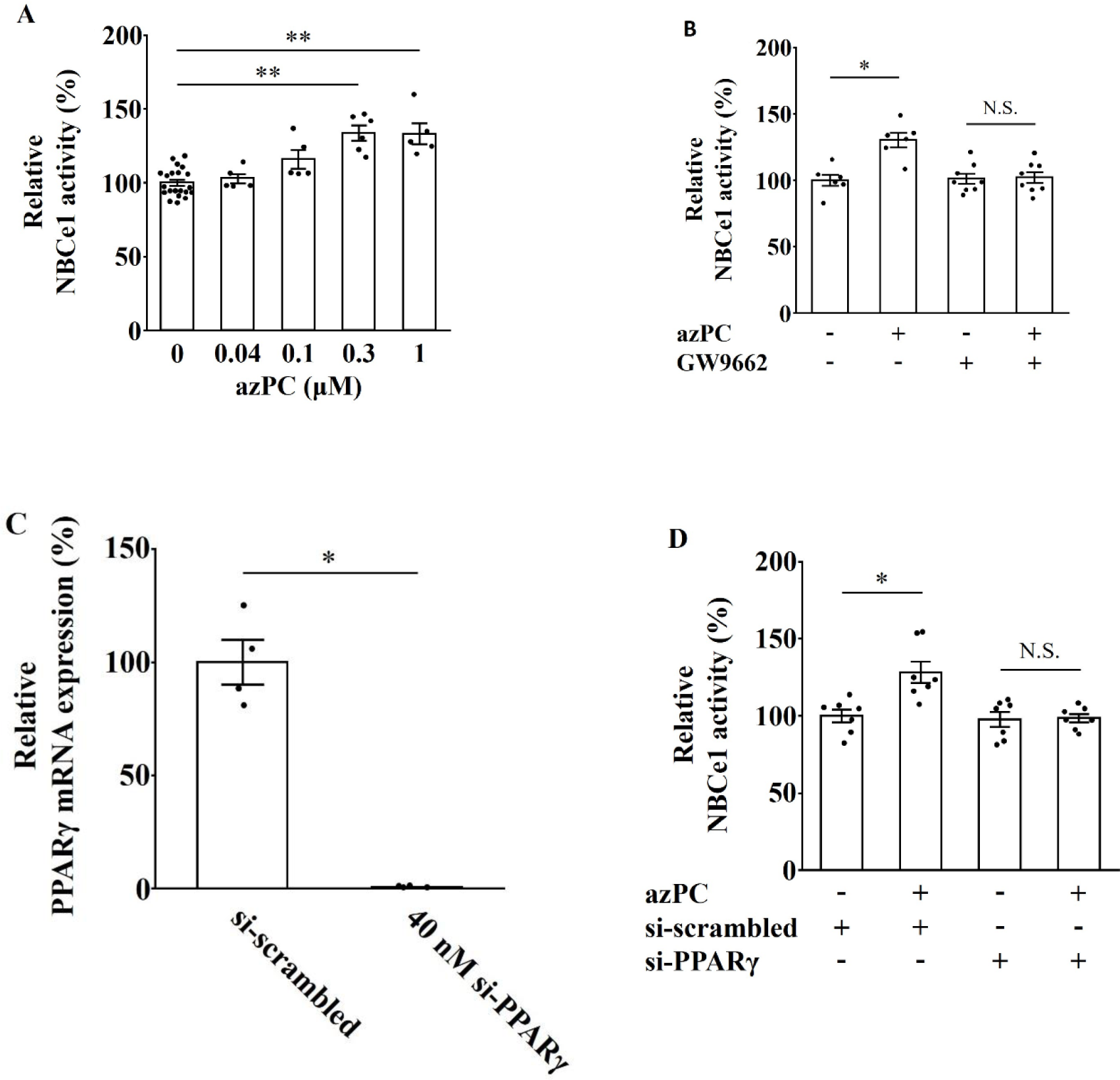
Effects of 1-O-hexadecyl-2-azelaoyl-*sn*-glycero-3-phosphocholine (azPC) on Na^+^/HCO_3_^-^ cotransporter 1 (NBCe1) activity in rat proximal tubules (PTs). (A) Effects of azPC in the concentration range from 0.04 to 1 μM on NBCe1 activity in isolated rat PTs. Control, n = 21; 0.04 μM azPC, n = 5; 0.1 μM azPC, n = 5; 0.3 μM azPC, n = 6; 1 μM azPC, n = 5; ** *p* < 0.01 versus control. (B) Effects of 5 μM GW9662 on NBCe1 activity in PTs treated with 0.3 μM azPC. Control, n = 6; azPC, n = 6; GW9662, n = 8; azPC + GW9662, n = 8; * *p* < 0.05 versus control. (C) Proliferator-activated receptor γ (PPARγ) mRNA expression in isolated rat PTs treated with siRNA against PPARγ at 40 nM (si-PPARγ) as compared to that in isolated rat PTs treated with scrambled negative control (si-scrambled). n = 4; * *p* < 0.05 versus si-scrambled. (D) Effects of siRNA treatment on 0.3 μM azPC-stimulated NBCe1 activity in isolated rat PTs. PTs were treated with si-scrambled or si-PPARγ. n = 7; * *p* < 0.05 versus si-scrambled. Each open bar represents the relative activity of NBCe1. NBCe1 activity of control group (azPC-untreated PT) was set at 100%.

We next examined whether the stimulation of NBCe1 activity by azPC was dependent on PPARγ signaling using a specific PPARγ antagonist, 2-chloro-5-nitro-N-phenylbenzamide (GW9662, 5 μM). GW9662 completely inhibited the stimulatory effects of azPC on NBCe1 activity without affecting the basal NBCe1 activity (Figure 1B). We also performed gene silencing experiments with siRNA against PPARγ in cultured rat PTs, as previously described (20, 31). As shown in Figure 1C, 40 nM siRNA against PPARγ significantly suppressed the expression of PPARγ mRNA as compared to the scrambled negative control. Additionally, treatment with 40 nM siRNA against PPARγ did not affect the basal NBCe1 activity as compared to the treatment with scrambled negative control, which completely blocked the stimulation of NBCe1 by azPC (Figure 1D). These results indicate that azPC stimulates NBCe1 activity via PPARγ-dependent signaling.

### Effects of azPC on luminal NHE activity in isolated rat PTs

Next, we focused on luminal NHE activity in freshly isolated rat PTs. Luminal NHE activity was measured using lumen-opened PTs, as previously described (22,31–33). The stimulatory effects of azPC on luminal NHE activity were observed in a dose-dependent manner (Figures 2A and S2), similar to NBCe1 activity. Therefore, we used 0.3 μM azPC for subsequent experiments.

**Figure 2.**
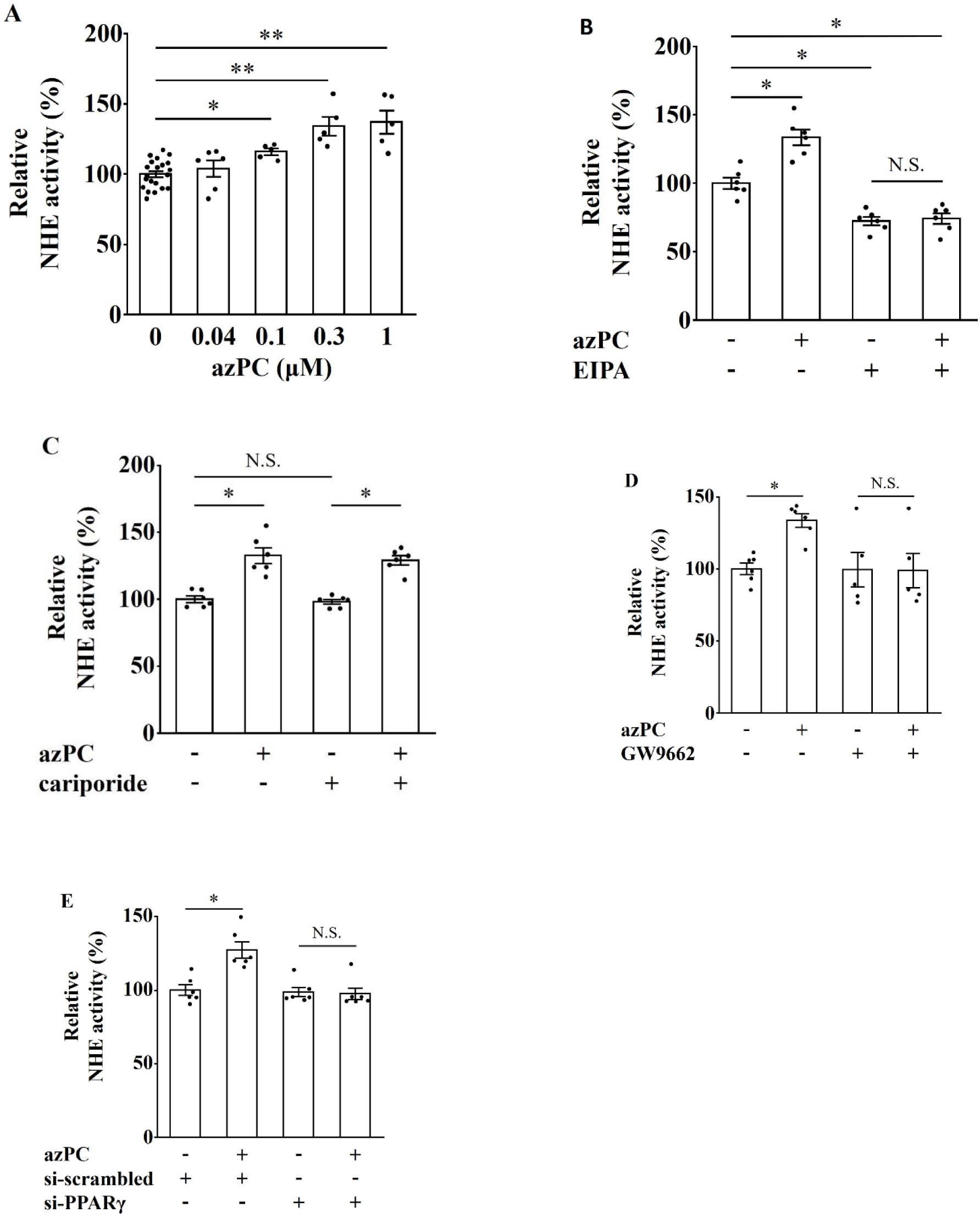
Effects of azPC on luminal Na^+^/H^+^ exchanger (NHE) activity in rat PTs. (A) Effects of azPC in the concentration range from 0.04 to 1 μM on luminal NHE activity in isolated rat PTs. Control, n = 21; 0.04 μM azPC, n = 6; 0.1 μM azPC, n = 5; 0.3 μM azPC, n = 5; 1 μM azPC, n = 5; * *p* < 0.05, ** *p* < 0.01 versus control. (B) Effects of 100 μM ethyl-isopropyl amiloride on luminal NHE activity in PTs treated with 0.3 μM azPC. n = 6; * *p* < 0.05 versus control. (C) Effects of 1 μM cariporide on luminal NHE activity in PTs treated with 0.3 μM azPC. n = 6; * *p* < 0.05 versus control. (D) Effects of 5 μM GW9662 on luminal NHE activity in PTs treated with 0.3 μM azPC. Control, n = 6; azPC, n = 6; GW9662, n = 5; azPC + GW9662, n = 5; * *p* < 0.05 versus control. (E) Effects of siRNA treatment on 0.3 μM azPC-stimulated luminal NHE activity in isolated rat PTs. PTs were treated with si-scrambled or siRNA against PPARγ at 40 nM. n = 6; * *p* < 0.05 versus si-scrambled. Each PT was also treated with 200 nM bafilomycin A_1_. Each open bar represents the relative activity of luminal NHEs. NHE activity of the control group (azPC-untreated PT) was set at 100%.

Next, we performed experiments using an NHE inhibitor, ethyl-isopropyl amiloride (EIPA, 100 μM). As shown in Figure 2B, the stimulatory effects of azPC were completely inhibited by EIPA. EIPA treatment partially, but significantly, decreased the basal activity by approximately 30% (Figure 2B). Next, we examined the effect of cariporide (1 μM) on NHE activation by azPC. Figure 2C shows that cariporide did not affect the basal activity or azPC-induced stimulatory responses. These results suggest that azPC stimulates luminal NHE activity in rat PTs.

We next investigated whether the stimulation of luminal NHE activity by azPC was mediated by PPARγ using GW9662 (5 μM). GW9662 completely inhibited the stimulatory effects of azPC on luminal NHE activity without affecting the basal NHE activity (Figure 2D). Furthermore, we performed gene-silencing experiments using siRNA against PPARγ at 40 nM. As shown in Figure 2E, siRNA treatment against PPARγ did not affect the basal NHE activity as compared to the treatment with scrambled negative control, which completely suppressed the stimulation of luminal NHE by azPC. Therefore, these results indicate that the stimulation of NHE activity by azPC is mediated by PPARγ.

### Signaling pathway for stimulation of NBCe1 and NHE activities by azPC in rats

We previously reported that TZDs stimulate PT sodium transport through the PPARγ/Src/EGFR/ERK pathway (22). In this study, we confirmed whether the signaling mechanism of azPC-induced PT transport stimulation overlaps with that of TZD-induced PT transport stimulation. We examined the effect of a mitogen-activated protein/extracellular signal-regulated kinase (MEK) inhibitor, PD98059 (10 μM), on azPC-induced stimulation of PT transport by measuring NBCe1 and luminal NHE activities in freshly isolated rat PTs, and we analyzed the impact of azPC on ERK phosphorylation in rat kidney cortex tissues using western blot analysis. As shown in Figures 3A and 3B, PD98059 did not affect the basal activities of NBCe1 and NHE, and it completely inhibited the stimulatory effects of azPC on both NBCe1 and NHE activities. Western blot analysis revealed that azPC induced ERK phosphorylation in a dose-dependent manner (Figures S3 and S4). Furthermore, azPC-induced ERK phosphorylation was completely blocked by GW9662 and PD98059 (Figures 3C-3F). These results suggest that azPC-induced stimulation of PT transport is dependent on the PPARγ/MEK/ERK signaling pathway.

**Figure 3.**
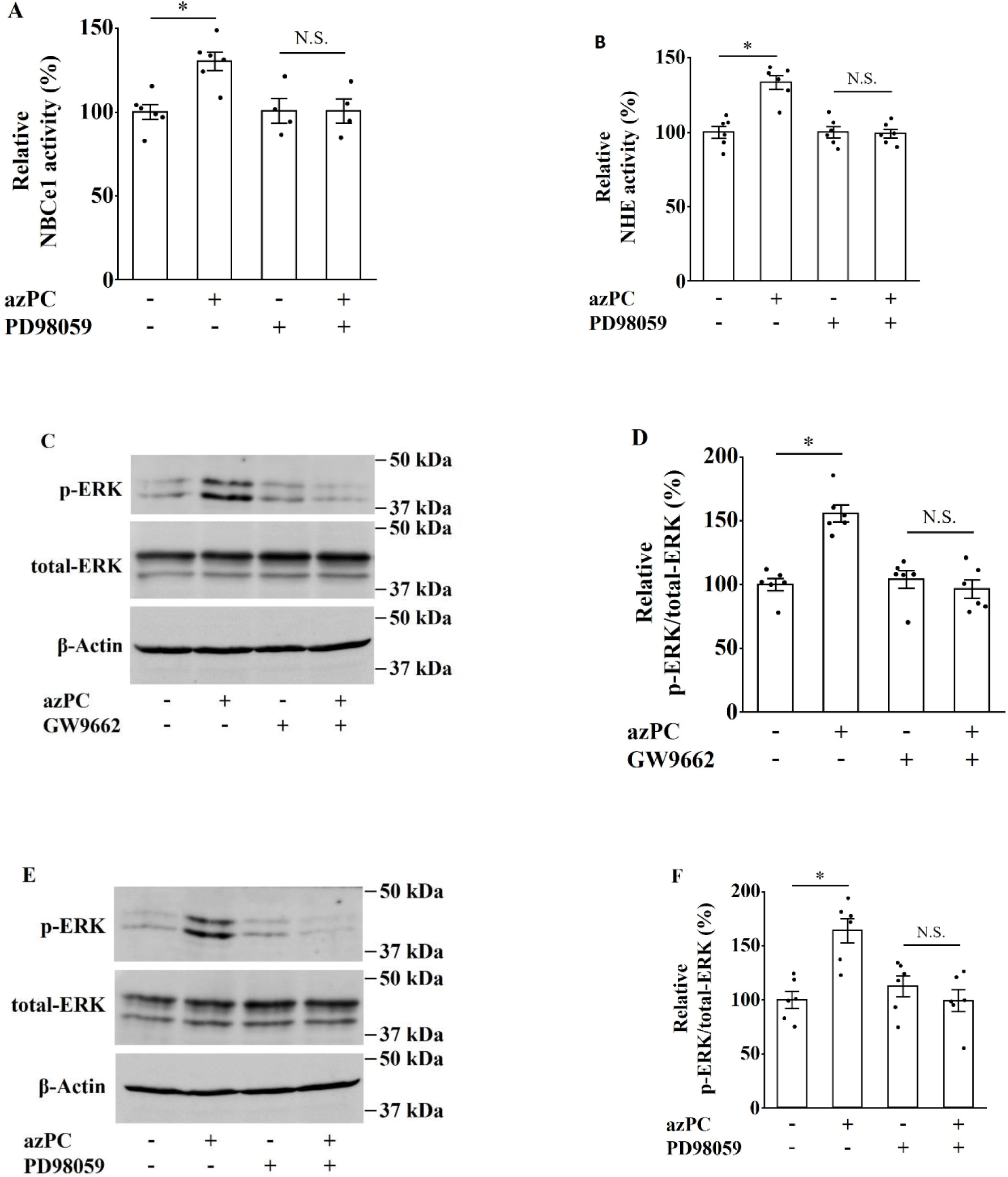
Role of extracellular signal-regulated kinase (ERK) in azPC-induced PPARγ-dependent pathway in rat PTs. (A) Effects of 10 μM PD98059 on NBCe1 activity in PTs treated with 0.3 μM azPC. Control, n = 6; azPC, n = 6; PD98059, n = 4; azPC + PD98059, n = 4; * *p* < 0.05 versus control. (B) Effects of 10 μM PD98059 on luminal NHE activity in PTs treated with 0.3 μM azPC. n = 6; * *p* < 0.05 versus control. (C) ERK phosphorylation in rat kidney cortex tissues. Kidney samples were treated with 0.3 μM azPC in the presence or absence of 5 μM GW9662. (D) Effects of 5 μM GW9662 on azPC-induced phosphorylation of ERK in rat kidney cortex tissues. n = 6; * *p* < 0.05 versus azPC-untreated and GW9662-untreated kidney cortex. (E) ERK phosphorylation in rat kidney cortex tissues. Kidney samples were treated with 0.3 μM azPC in the presence or absence of 10 μM PD98059. (F) Effects of 10 μM PD98059 on azPC-induced phosphorylation of ERK in rat kidney cortex tissues. n = 6; * *p* < 0.05 versus azPC-untreated and PD98059-untreated kidney cortex.

### Effects of azPC on PT transport in humans

Next, we examined the effect of azPC on PT transport in humans. No patient showed a severe renal dysfunction (Table S1). Addition of 0.3 μM azPC stimulated both NBCe1 and luminal NHE activities in freshly isolated human PTs, and the stimulatory responses were completely suppressed by GW9662 (Figures 4A and 4B). Moreover, western blot analysis in human kidney cortex tissues revealed that 0.3 μM azPC significantly enhanced ERK phosphorylation, and the enhancement of ERK phosphorylation was completely blocked by GW9662 (Figures 4C and 4D). Thus, we observed that azPC stimulated human PT transport through the PPARγ/MEK/ERK signaling pathway as well as rat PT transport.

**Figure 4.**
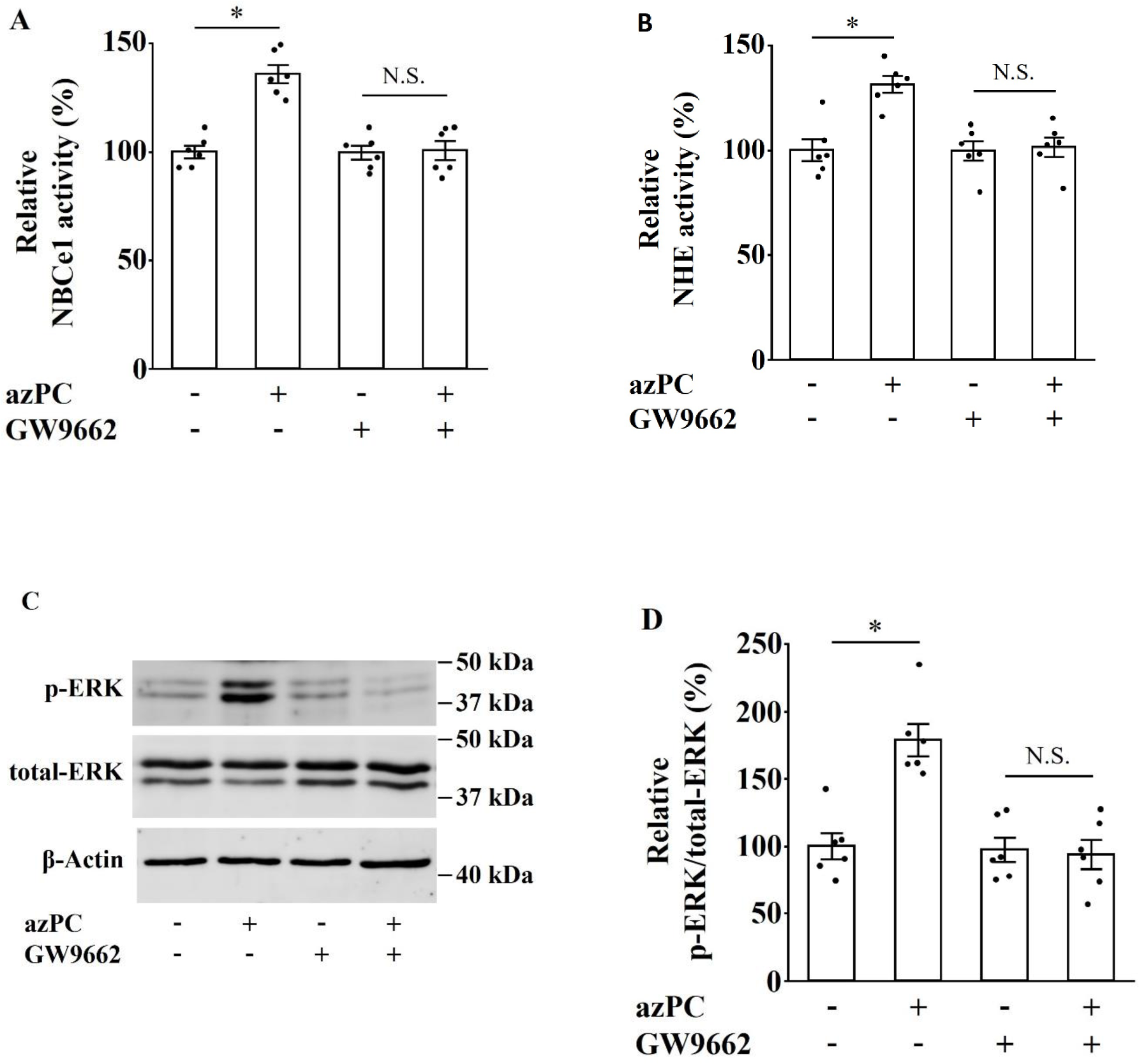
PPARγ-dependent stimulation of NBCe1 and luminal NHE activities by azPC in humans. (A) Effects of 5 μM GW9662 on NBCe1 activity in human PTs treated with 0.3 μM azPC. n = 6; * *p* < 0.05 versus control. (B) Effects of 5 μM GW9662 on luminal NHE activity in human PTs treated with 0.3 μM azPC. n = 6; * *p* < 0.05 versus control. (C) ERK phosphorylation in human kidney cortex tissues. Kidney samples were treated with 0.3 μM azPC in the presence or absence of 5 μM GW9662. (D) Effects of 5 μM GW9662 on azPC-induced phosphorylation of ERK in human kidney cortex tissues. n = 6; * *p* < 0.05 versus azPC-untreated and GW9662-untreated kidney cortex.

## Discussion

In this study, we demonstrated that azPC rapidly stimulated renal PT sodium transport by activating both NBCe1 and luminal NHEs in rats and humans, and the stimulatory responses were mediated by PPARγ. The azPC-induced activation of NBCe1 and NHE was inhibited by PD98059 in isolated rat PTs. Additionally, azPC enhanced ERK phosphorylation in kidney cortex tissues, and azPC-induced ERK phosphorylation was inhibited by GW9662 and PD98059. These results suggest that azPC stimulates sodium reabsorption from rat and human PTs through the PPARγ/MEK/ERK signaling pathway (Figure 5).

**Figure 5.**
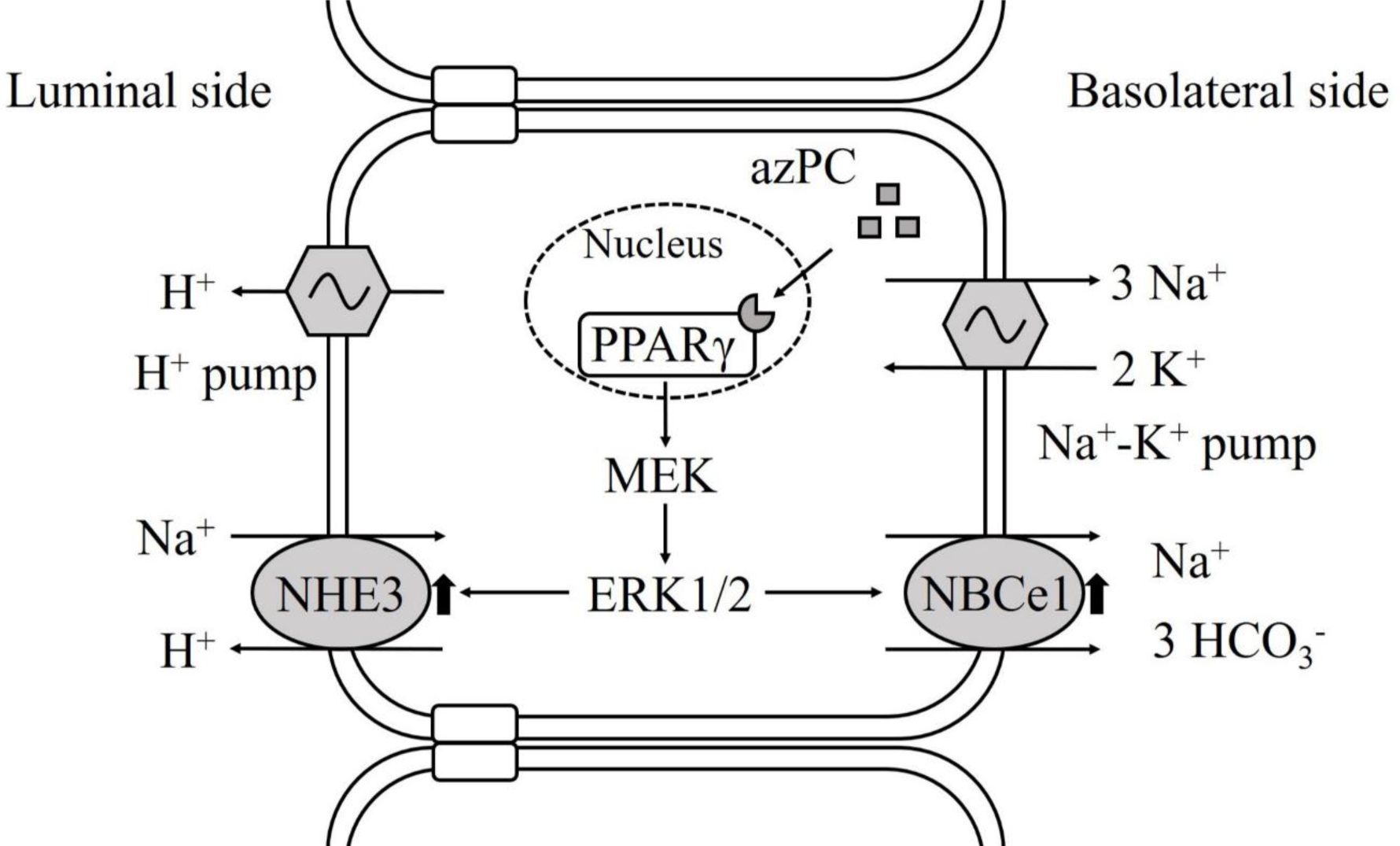
Schematic representation of the effect of azPC on sodium transport in PTs. azPC activates basolateral NBCe1 and luminal NHE3 through the PPARγ/MEK/ERK signaling pathway, leading to the stimulation of sodium transport in PTs.

The action of TZDs on sodium transporters in PT has been reported previously (34–36). For example, troglitazone has been reported to activate NBCe1 activity in rabbit PTs (34). Other studies have also reported that TZDs enhance the expression of NHE3 in human proximal tubule cells and rat kidneys (35, 36). However, the effect of endogenous PPARγ ligands on sodium transporters in PT is unknown. In this study, we demonstrated that 0.3 µM azPC stimulated NBCe1 activity by approximately 34 ± 5% in isolated rat PTs. Based on our previous findings that TZDs stimulate NBCe1 activity by approximately 35%-40% in rat PTs (22), the degree of NBCe1 activation by azPC is almost comparable to that of NBCe1 activation by TZDs. Therefore, the effect of azPC on PT transport seems likely to be involved in volume expansion, although the effect of azPC on other nephrons, including collecting ducts, needs to be elucidated.

We measured NHE activity by calculating the rate of decrease of intracellular pH (pH_i_) caused by bath Na^+^ removal and buffer capacity using lumen-opened PTs (31). To confirm the activation of NHE by azPC, we demonstrated that azPC-induced stimulation was completely inhibited by 100 µM EIPA (Figure 2B). Furthermore, EIPA decreased the basal activity by approximately 30% (Figure 2B), which seemed less effective than other studies because several previous studies have reported that EIPA reduces HCO_3_^-^ reabsorption in PTs by 40%-60% using a microperfusion technique (32,37,38). Our results are presumed to be affected by sodium-coupled transporters other than luminal NHEs. Several studies have suggested the involvement of transporters other than NHE3 and V-ATPase in luminal sodium-coupled bicarbonate absorption in PTs (32,37–39). The presence of a novel NBC on the luminal membranes of PTs has also been proposed in a recent study (40). Sodium-coupled transporters expressed on the luminal membranes of PTs can be reflected in our method while the function of NBCe1 is not significantly affected by Na^+^ concentration (41). Cariporide was used to confirm whether azPC activated NHE1 expressed on the basolateral membranes of PTs (Figure 2C) (42). The results showed that cariporide did not affect NHE activation by azPC, indicating that azPC activated luminal NHEs. Moreover, comparative studies using a microperfusion technique showed that the contribution of NHE2 to Na^+^-H^+^ translocation in PTs was less than that of NHE3 (37, 43). Based on these results, we determined that azPC can activate NHE3.

Nuclear receptors, including PPARγ, have been reported to mediate both non-genomic and genomic actions (44). The short time frame in the range of seconds to minutes is essential to distinguish between nongenomic and genomic actions (44). The azPC-induced actions on PTs were exerted rapidly within a few minutes, which were consistent with the features of nongenomic actions. Furthermore, we found that azPC activated luminal NHEs in rat and human PTs but not in mouse PTs (Figure S5), which was similar to TZD-induced actions (22). Therefore, these findings suggest that the PPARγ-dependent signaling pathway activated by azPC may overlap with the nongenomic signaling pathway activated by TZDs. Although a variety of nongenomic signaling activated by PPARγ ligands has been described (45), the MEK/ERK pathway probably plays an important role in PT transport because multiple ligands such as TZDs and Ang II have been reported to activate both NBCe1 and NHE3 through the MEK/ERK pathway (19, 22).

The stimulatory effects of azPC on PT sodium transport were dose-dependent at submicromolar concentrations. The pharmacokinetics of azPC have not been fully understood, and the physiological concentration of azPC in PT is unknown. However, the concentrations of azPC used in the present study should be reasonable. It has been demonstrated that azPC dose-dependently induces peroxisome proliferator-activated receptor response element (PPRE) reporter gene expression at submicromolar concentrations in CV-1 cells transfected with acyl-CoA-oxidase-PPRE-luciferase reporter plasmid (30). Other studies have also reported that 1 µM azPC exhibits sufficient PPARγ activation comparable to that of TZDs (46, 47). Additionally, the plasma of humans and rodents contains low micromolar levels of total oxidatively fragmented PCs, and the plasma concentrations of several types of fragmented PCs such as 1-palmitoyl-2-(5-oxovaleroyl)-*sn*-glycero-3-phosphocholine are in low micromolar or submicromolar ranges. Thus, previous studies likely support the validity of the concentrations of azPC in the present study (7,11,48).

Cluster determinant 36 (CD36) is a multifunctional receptor that mediates the cellular uptake of a variety of oxidation products, including oxPLs and oxLDL, and it is widely expressed in PTs of the kidney (49). Regarding the relationship between CD36 and azPC, a previous study has reported that in human monocytes, azPC enhances CD36 expression and CD36 promotes the uptake of extracellular azPC (30). Although the association between azPC and CD36 in PT transport remains unknown, CD36 might promote the uptake of azPC in PT because it has been suggested that CD36 mainly mediates the internalization of oxLDL in tubular cells (50). Additionally, other studies have demonstrated that the downstream signaling triggered by renal CD36 includes the MEK/ERK pathway in human renal proximal tubule epithelial cells and human embryonic kidney 293 cell lines (51, 52). These findings suggest an overlap between PPARγ/MEK/ERK signaling stimulated by azPC and CD36 signaling in PT. Whether the stimulatory effects of azPC on PT transport were mediated by CD36 was not investigated in the present study because we could not obtain a CD36 antagonist, such as apolipoprotein A-I-mimetic peptide 5A (53), which is not commercially available. Although gene silencing experiments with siRNA against CD36 were performed in cultured rat PTs, the expression of CD36 mRNA was not significantly suppressed as compared to that of the scrambled negative control (data not shown).

In summary, we demonstrated that azPC rapidly activated basolateral NBCe1 and luminal NHEs via the PPARγ/MEK/ERK pathway in isolated PTs from rat and human kidneys. The stimulation of PT sodium and water reabsorption by azPC is likely a novel mechanism leading to the development of volume expansion. We believe that these findings can provide an impetus for elucidating the mechanism of atherosclerosis-induced volume expansion and hypertension because azPC is strongly associated with the development and progression of atherosclerosis.

## Experimental procedures

### Animal studies

Male Wistar rats and male C57BL/6 mice were purchased from CLEA Japan, Inc. (Tokyo, Japan). They were housed in cages with a 12/12 h light/dark cycle, and they were provided standard food and water *ad libitum*. Rats and mice, at 4–6 weeks of age, were sacrificed after anesthetization with excessive amounts of pentobarbital sodium (Somnopentyl; Kyoritsu Seiyaku, Tokyo, Japan) (intraperitoneally, 50 mg/kg), and the kidney samples were obtained. All animal experiments were performed in accordance with local institutional guidelines (authorization number: P17-070).

### Human samples

Human kidney samples were obtained from patients who underwent unilateral nephrectomy for renal carcinoma. The study was approved by the Institutional Review Board of the University of Tokyo School of Medicine (2520-[11]), and signed informed consent was obtained from all subjects.

### Measurements of NBCe1 activity in renal PTs from rats and humans

NBCe1 activity was determined as previously described (18,22,54). Briefly, the PT (S2 segment) fragment was manually microdissected from rat or human kidneys without collagenase treatment, and it was transferred to a perfusion chamber mounted on an inverted microscope. To avoid the influence of luminal transporters, the PT fragment was collapsed with two holding pipettes. The luminally collapsed PT was incubated with an acetoxymethyl ester form of a pH-sensitive fluorescent dye 2’,7’-bis(carboxyethyl)-5(6)-carboxyfluorescein acetoxymethyl ester (BCECF/AM; Dojindo Laboratories, Kumamoto, Japan) in Dulbecco’s modified Eagle’s medium (DMEM) for 10 min, and pH_i_ was monitored with a photometry system, MetaFluor 7.7 software (Molecular Devices, Sunnyvale, CA, USA). The chamber was perfused with prewarmed (38 °C) DMEM equilibrated with 5% CO_2_/95% O_2_ gas, and subsequently, bath HCO_3_^-^ concentrations were repeatedly switched from 25 mM to 12.5 mM in the absence and presence of azPC (Cayman Chemical, Ann Arbor, MI, USA) or other chemical agents such as a specific PPARγ antagonist GW9662 (Sigma-Aldrich, St Louis, MO, USA) at 5 µM and an MEK inhibitor 2-(2-amino-3methoxyphenyl)chromone (PD98059) (FUJIFILM Wako Pure Chemical, Osaka, Japan) at 10 µM, both of which exhibit sufficient inhibitory activities without affecting the basal NBCe1 activity in PTs (19, 22). NBCe1 activity was calculated using the rate of pH_i_ decrease in response to bath HCO_3_^-^ reduction and buffer capacity.

### Measurements of luminal NHE activity in renal PTs from rats, mice, and humans

Luminal NHE activity was determined as previously described (22,31,33). Briefly, the PT (S2 segment) fragment was freshly isolated in the same way as the measurements of NBCe1 activity, and they were attached to a glass coverslip with Cell-tak glue (Corning, One Riverfront Plaza, NY, USA). The tubule was placed on a perfusion chamber mounted on an inverted microscope, and the end of the tubule was cut with a capillary glass to sufficiently expose the lumen of the tubule. The lumen-opened PT was incubated with BCECF/AM in HEPES-buffered solution (144 mM Na^+^, 5 mM K^+^, 1.5 mM Ca^2+^, 1 mM Mg^2+^, 137 mM Cl^-^, 2 mM H_2_PO_4_^-^, 1 mM SO_4_, 5.5 mM glucose, 25 mM HEPES, adjusted to pH 7.4) (31, 55) for 10 minutes, and pH_i_ was monitored with MetaFluor 7.7 software. A pre-warmed (38 °C) HEPES-buffered solution was used for the bath perfusate, and 200 nM bafilomycin A_1_ (FUJIFILM Wako Pure Chemical) was added to block the effect of V-ATPase on PT transport (31, 33). The perfusate was repeatedly switched from HEPES-buffered solution to an isotonic Na^+^-free solution (144 mM N-methyl-D-glucamine, 5 mM K^+^, 1.5 mM Ca^2+^, 1 mM Mg^2+^, 137 mM Cl^-^, 2 mM H_2_PO_4_^-^, 1 mM SO_4_^2-^, 5.5 mM glucose, 25 mM HEPES, adjusted to pH 7.4) in the absence and presence of azPC or other chemical agents such as two NHE inhibitors EIPA (Research Biochemicals Incorporated, Natick, MA, USA) at 100 µM and cariporide (Santa Cruz Biotechnology, Dallas, TX, USA) at 1 µM and GW9662 at 5 µM and PD98059 at 10 µM. EIPA was used at a concentration that significantly inhibited all isoforms of NHE (32, 56). Cariporide was used at a concentration that significantly inhibited NHE1 but not NHE3 in murine cells (56–58). GW9662 (5 µM) and PD98059 (10 µM) exhibited sufficient inhibitory activities without affecting the basal NHE activity in PTs (19, 22). Luminal NHE activity was calculated using the rate of pH_i_ decrease caused by bath Na^+^ removal and buffer capacity.

### siRNA treatment in isolated rat PTs

siRNA treatment of isolated rat PTs was performed as previously described (20, 31). Briefly, freshly isolated rat PTs were treated with siRNA against PPARγ (AM16708; Invitrogen, Carlsbad, CA, USA) at 40 nM, siRNA against CD36 (4390771; Invitrogen), or scrambled negative control (sc-37007; Santa Cruz Biotechnology) using Lipofectamine 2000 and Opti-MEM I Reduced Serum Medium (both from Invitrogen). The PTs were incubated in DMEM supplemented with 10% fetal bovine serum at 37 °C overnight, and they were used to measure NBCe1 activity, luminal NHE activity, and quantitative PCR.

### RNA extraction and quantitative PCR analysis

Total RNA was extracted from isolated rat PTs with isogen II (Nippon Gene, Tokyo, Japan), according to manufacturer’s instructions, and first-strand cDNA was synthesized using a cDNA Synthesis Kit (Takara, Tokyo, Japan), as previously reported (20). The mRNA expression levels were estimated using quantitative PCR (Prism 7000; Applied Biosystems, Foster City, CA, USA) with TaqMan Gene Expression Master Mix (Applied Biosystems) and TaqMan Gene Expression Assay kits, Rn00440945_m1 for rat PPARγ, Rn00580728_m1 for rat CD36, or Rn00667869_m1 for rat β-actin (Applied Biosystems). The mRNA levels were normalized to β-actin expression levels.

### Western blot analysis

Thin slices of kidney cortex were obtained from rats or humans, and they were divided into small bundles, as previously described (22, 59). The kidney samples were incubated in DMEM at 37 °C under 5% CO_2_ for 40 min in the presence or absence of inhibitors such as 5 µM GW9662 and 10 µM PD98059, and they were incubated for 15 min in DMEM containing 0.3 µM azPC. After incubation, the samples were homogenized in ice-cold buffer A (25 mM Tris-HCl [pH 7.4], 10 mM sodium orthovanadate, 10 mM sodium pyrophosphate, 100 mM sodium fluoride, 10 mM EDTA, 10 mM EGTA, and 1 mM phenylmethylsulfonyl fluoride) (22), and they were centrifuged at 12000 × g for 10 min. The supernatant from each sample was collected and divided into aliquots containing equal amounts (approximately 20 µg) of proteins. The samples were separated using 10% SDS-PAGE, and they were transferred onto nitrocellulose membranes. After the membranes were blocked with 5% skim milk in Tris-buffered saline (137 mM NaCl, 2.68 mM KCl, 25 mM Tris, adjusted to pH 7.4), they were incubated with primary antibodies at 4 °C overnight, and following this, they were incubated with horseradish peroxidase (HRP)-conjugated secondary antibodies at room temperature for 1 h. Primary antibodies against ERK1/2 (9102), phospho-ERK1/2 (Thr202/Tyr204) (9101), and β-actin (4970) were purchased from Cell Signaling Technology (Danvers, MA, USA). HRP-conjugated anti-rabbit IgG antibody (111-035-003) was purchased from Jackson ImmunoResearch Laboratories (West Grove, PA, USA). The protein bands were detected using a chemiluminescence detection system (ImageQuant LAS 4000 mini; GE Healthcare, Little Chalfont, UK).

### Statistical analysis

All data are expressed as the mean ± standard error of the mean. The data were analyzed with JMP Pro 14 software (SAS Institute, Cary, NC, USA) using a Wilcoxon signed-rank test or Kruskal–Wallis test followed by a Steel test or Steel–Dwass test, as appropriate. Statistical significance was set at a *p* value < 0.05.

## Data availability

The data supporting the findings of this study are included within the manuscript and its supporting information.

## Acknowledgements

We are grateful to Dr. Masashi Suzuki for helpful discussions on this work.

## Author contributions

N. S., S. H. and M. Nakamura designed the study. T. M., H. T. and M. Nakamura performed the experiments. Y. S. and H. K. collected human kidney samples. T. M. and M. Nakamura wrote the manuscript. M. Nangaku and M. Nakamura supervised the project. All authors reviewed and approved the final manuscripts.

## Funding and additional information

The study was supported by Daiichi Sankyo Co., Ltd. (Grant Number A20-1972) and Japan Society for the Promotion of Science (JSPS) KAKENHI (Grant Number 19K08671 and 19K17698).

## Conflict of interest

M. Nakamura has received honorarium from Kyowa Kirin Co., Ltd. and Daiichi Sankyo Co., Ltd. All other authors declare that they have no conflicts of interest with the contents of this article.

## Abbreviations^1^

^1^oxidized phospholipids (oxPLs)

ethyl-isopropyl amiloride (EIPA)

oxidized low-density lipoproteins (oxLDLs)

peroxisome proliferator-activated receptor response element (PPRE)

1-O-hexadecyl-2-azelaoyl-*sn*-glycero-3-phosphocholine (azPC)

peroxisome proliferator-activated receptor γ (PPARγ)

vacuolar-type H^+^-ATPase (V-ATPase)

Na^+^/HCO_3_^-^ cotransporter 1 (NBCe1)

2’,7’-bis(carboxyethyl)-5(6)-carboxyfluorescein acetoxymethyl ester (BCECF/AM)

Dulbecco’s modified Eagle’s medium (DMEM)

angiotensin II (Ang II)

Intracellular pH (pH_i_)

horseradish peroxidase (HRP)

mitogen-activated protein/extracellular signal-regulated kinase (MEK)

Na^+^/H^+^ exchanger 3 (NHE3)

thiazolidinediones (TZDs)

Cluster determinant 36 (CD36)

“This article contains supporting information.” Any references cited in the Supporting Information should be cited in this sentence.

## Supporting Information

**Figure S1.**
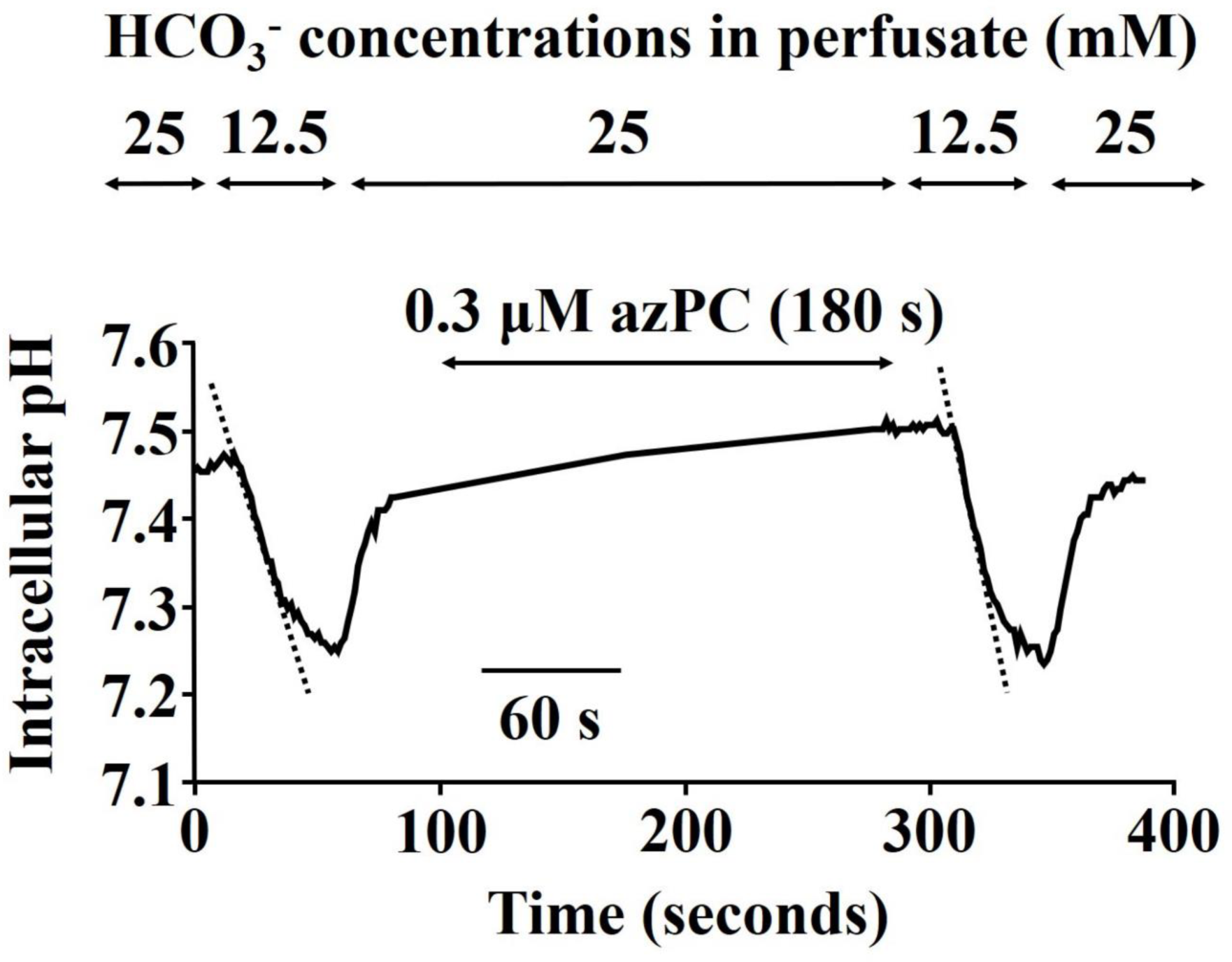
Intracellular pH (pH_i_) decrease in response to bath HCO_3_ reduction from 25 mM to 12.5 mM in isolated rat proximal tubules (PTs). PT exhibited a rapid decrease in pH_i_ 3 min after the addition of 0.3 µM azPC.

**Figure S2.**
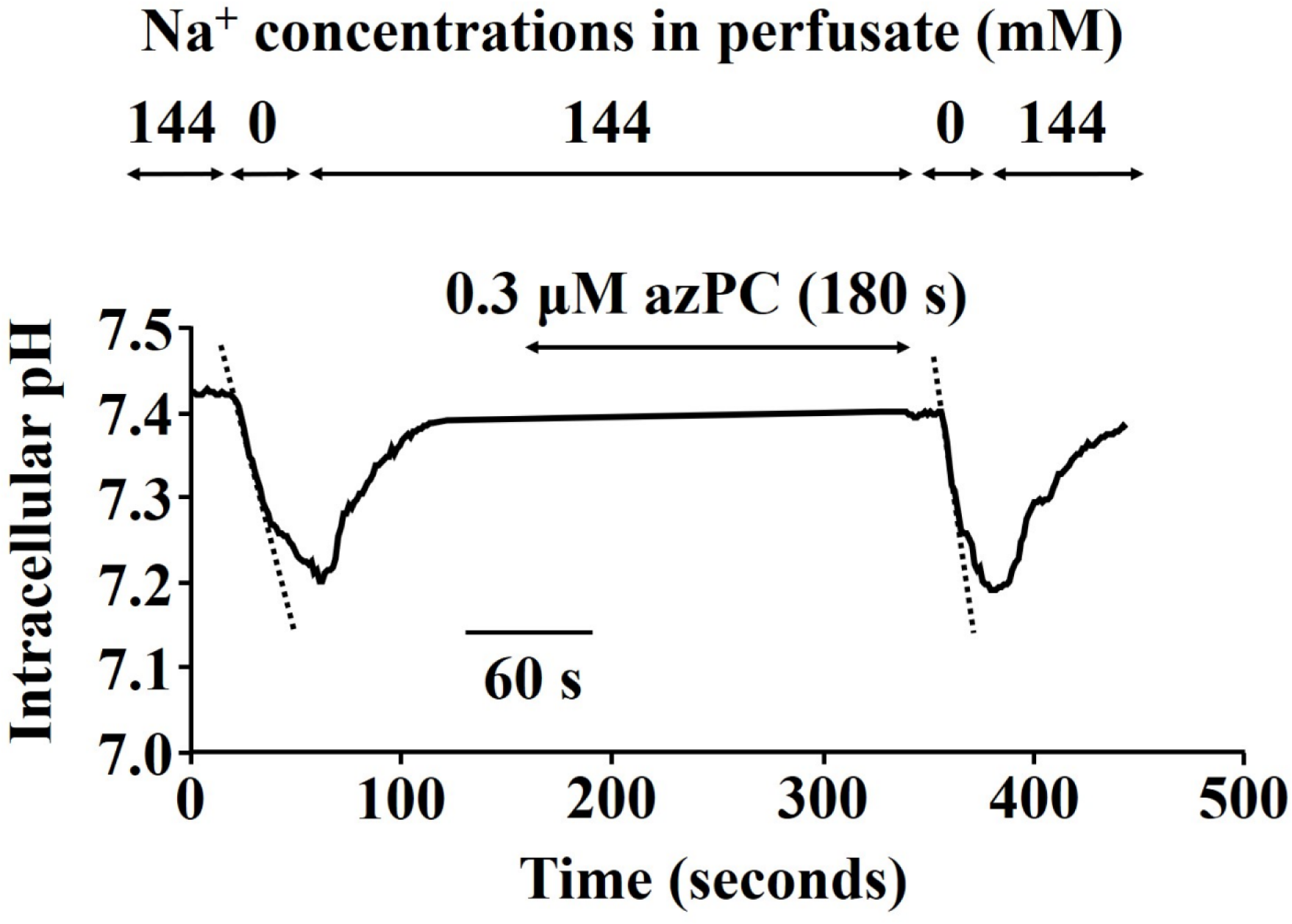
pH_i_ decrease in response to bath Na^+^ reduction from 144 mM to 0 in isolated rat PTs. Bafilomycin A_1_ (200 nM) was added to the perfusate to block the effect of V-ATPase. PT exhibited a rapid pH_i_ decrease after 3 min of addition of 0.3 µM azPC.

**Figure S3.**
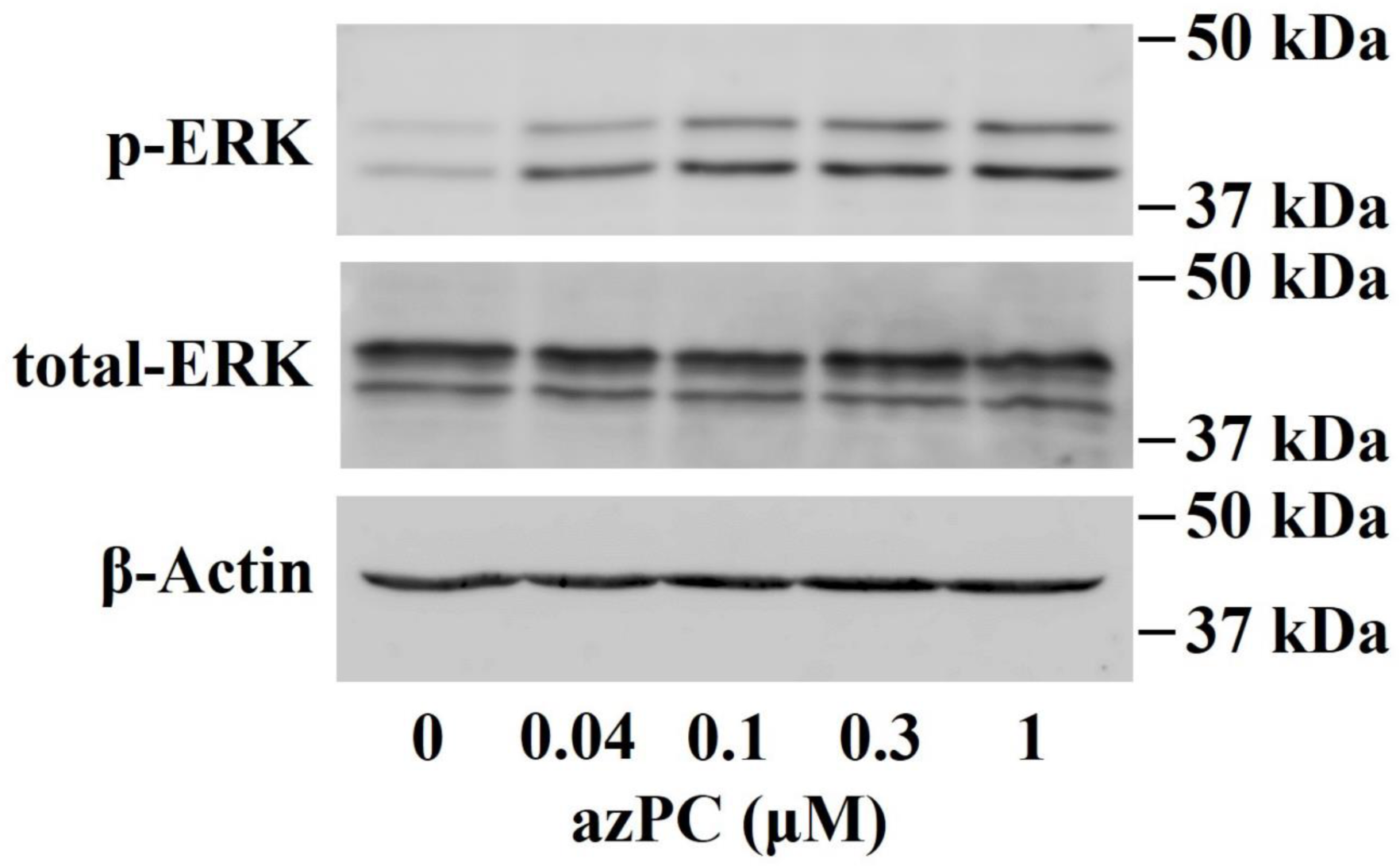
ERK phosphorylation in rat kidney cortex tissues. Kidney samples were treated with azPC at concentrations from 0.04 μM to 1 μM.

**Figure S4.**
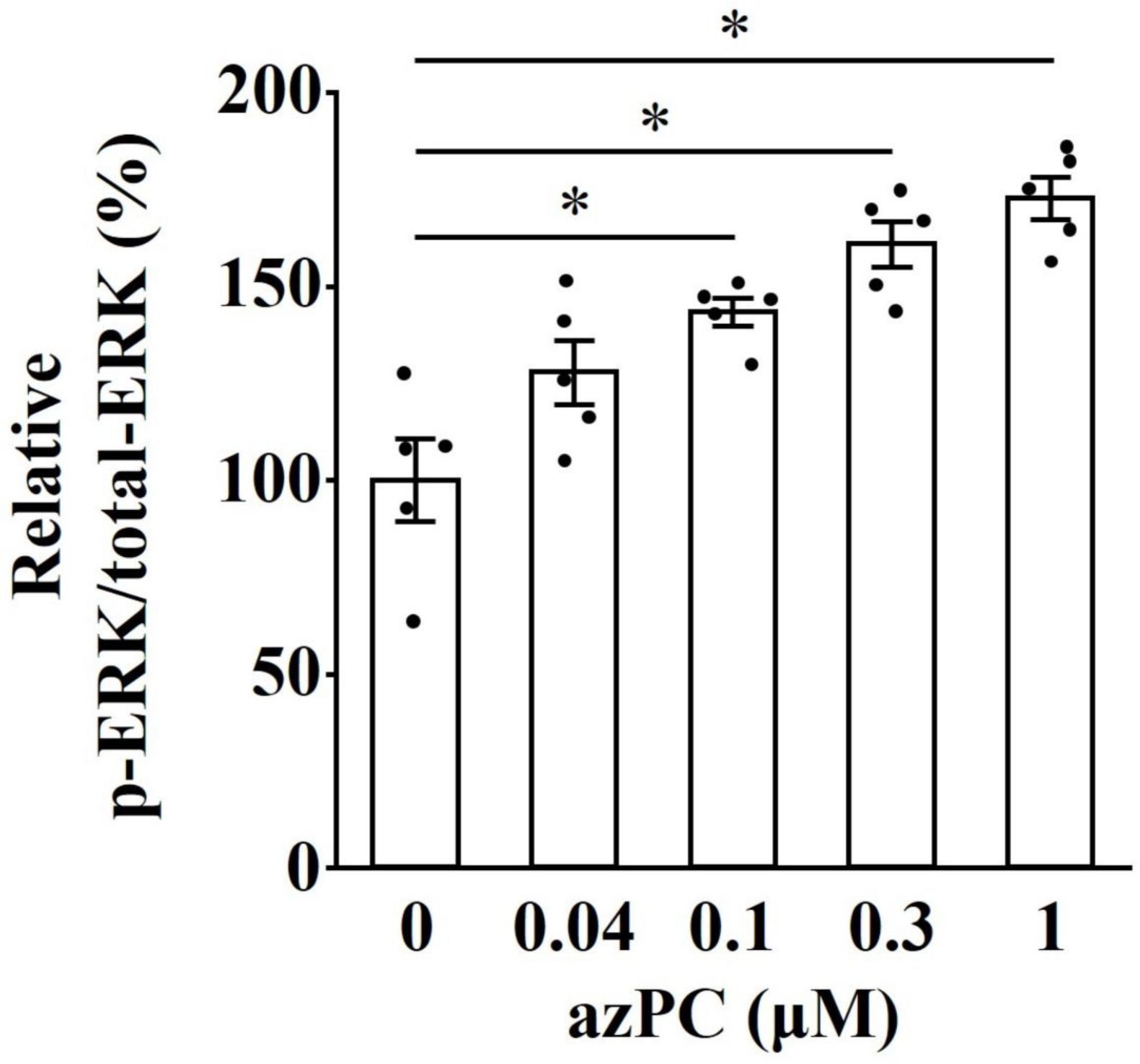
Effects of azPC at concentrations from 0.04 μM to 1 μM on ERK phosphorylation in isolated rat PTs. n = 5; * *p* < 0.05 versus azPC-untreated kidney cortex.

**Figure S5.**
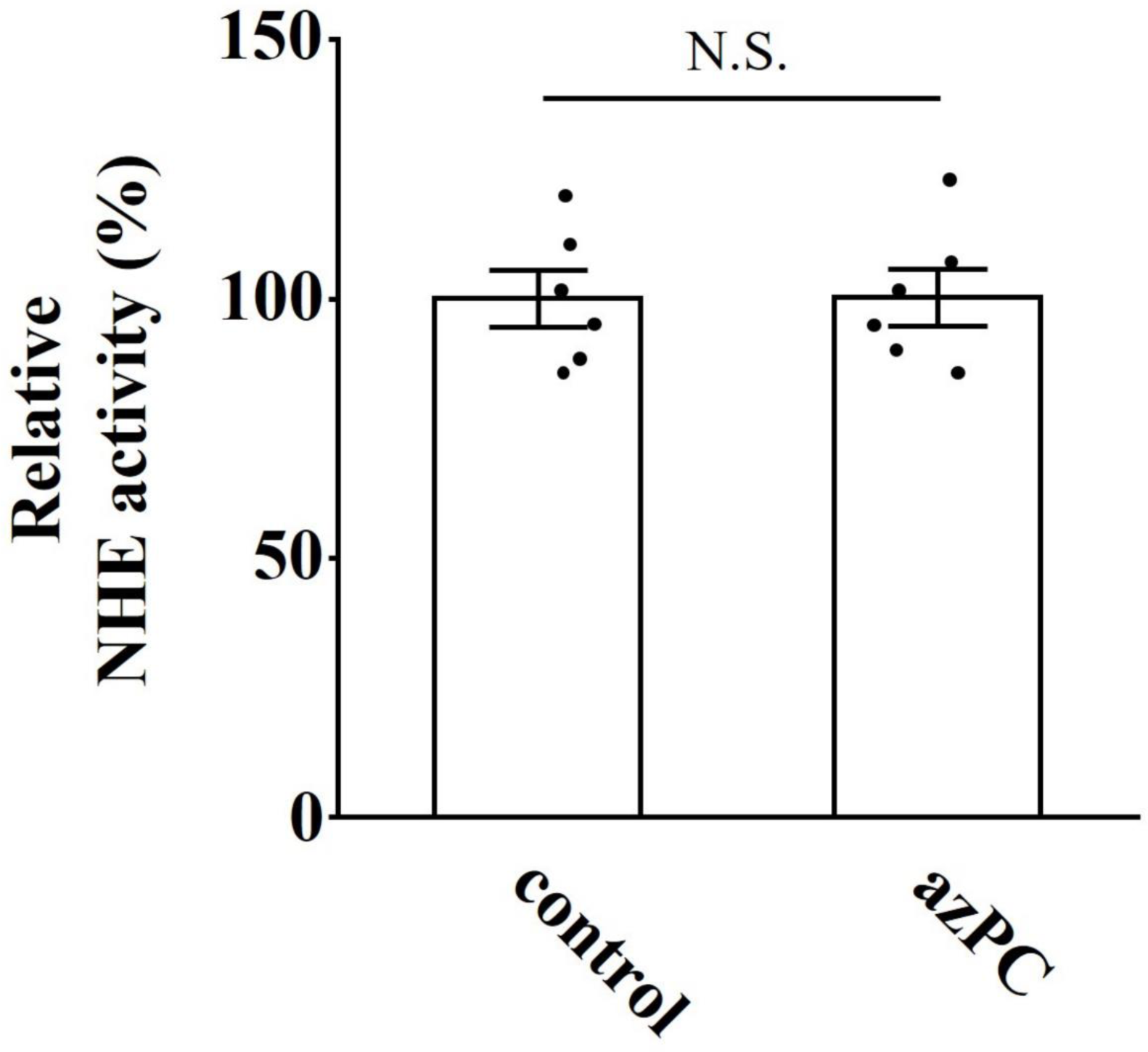
Effects of 0.3 μM azPC on luminal NHE activity in isolated mouse PTs. Each PT was treated with 200 nM bafilomycin A_1_. Each open bar represents the relative activity of luminal NHEs. NHE activity of the control group was set as 100%. n = 6.

